# Effect of imputation on gene network reconstruction from single-cell RNA-seq data

**DOI:** 10.1101/2021.04.13.439623

**Authors:** Lam-Ha Ly, Martin Vingron

## Abstract

Despite the advances in single-cell transcriptomics the reconstruction of gene regulatory networks remains challenging. Both the large amount of zero counts in experimental data and the lack of a consensus preprocessing pipeline for single-cell RNA-seq data make it hard to infer networks from transcriptome data. Data imputation can be applied in order to enhance gene-gene correlations and facilitate downstream data analysis. However, it is unclear what consequences imputation methods have on the reconstruction of gene regulatory networks.

To study this question, we evaluate the effect of imputation methods on the performance and structure of the reconstructed networks in different experimental single-cell RNA-seq data sets. We use state-of-the-art algorithms for both imputation and network reconstruction and evaluate the difference in results before and after imputation. We observe an inflation of gene-gene correlations that affects the predicted network structures and may decrease the performance of network reconstruction in general. Yet, within the modest limits of achievable results, we also make a recommendation as to an advisable combination of algorithms, while warning against the indiscriminate use of imputation before network reconstruction in general.

## 1 Introduction

Single-cell transcriptomics has revolutionized genomics. In particular, this new type of data is widely assumed to advance the unraveling of regulatory interactions in the cell. Thus, there is great interest in the computational reconstruction of gene regulatory networks (GRNs) from single-cell transcriptome data.

Available methods for GRN reconstruction from single-cell RNA-seq (scRNAseq) data draw on a plethora of statistical approaches (1–6)). Pratapa et. al. (6) provide an extensive benchmark study evaluating the performance of various methods. However, for GRN reconstruction several authors have remarked that preprocessing the data is important, mostly due to the sparse nature of the data (7,8). Several computational analysis pipelines have been suggested and are in wide use (9,10). Typically, as one of the early steps, such a pipeline will include a data normalization and/or imputation step, which statistically estimates unobserved read counts in cases where the method deems that experimental or technical noise has led to the absence of a count, i.e., a so-called dropout. While normalization attempts to correct for different read depths between cells (11,12), imputation attempts to recover gene counts by predicting missing data and eventually smoothen gene expression values (13–22). In some tools a prior normalization step is not required but integrated within the imputation method (20,21). Hou et. al (23) extensively evaluated the impact of imputation on clustering, differential expression analysis and pseudotime inference and invoked cautious interpretations of the results.

It still remains unclear though how imputation methods affect network structures (24). On the one hand, it is recommended to use imputation to enhance gene regulatory correlations prior to network inference (18,20). But on the other hand, results based on imputed data should be interpreted with care (10,23,25). Thus, imputation meets conflicting attitudes within the community.

Here, we systematically study the question whether data imputation as a preprocessing step affects results obtained using reconstructed GRNs. We build on previously published benchmark studies and consider the best-performing scRNAseq-based tools for both imputation and network reconstruction in our analysis. We measure the performance of different combinations of imputation method and GRN reconstruction method using multiple experimental datasets and a ground truth network that has been used in other benchmark studies. We compare the performance and network structures obtained using unimputed data and imputed data, respectively, and show that in most cases GRN reconstruction does not profit from imputation. In order to explain the observed results we analyze the effect of imputation on predicted gene interactions. Ultimately, we present a recommendation, how to proceed in a data analysis project.

## 2 Results

### 2.1 Systematic evaluation of network models

Evaluating the combination between imputation and network inference on different datasets results in a cubic matrix. To manage this we restrict our selection to state-of-the-art computational tools, both for imputation and network inference, that perform most accurately and have been recommended in recent benchmark studies (6,23). Consequently, we developed a computational pipeline to study seven cell types that were obtained from different scRNAseq experiments, using four state-of-the-art imputation methods combined with three top performing GRN methods as depicted in Figure 1.

**Figure 1.**
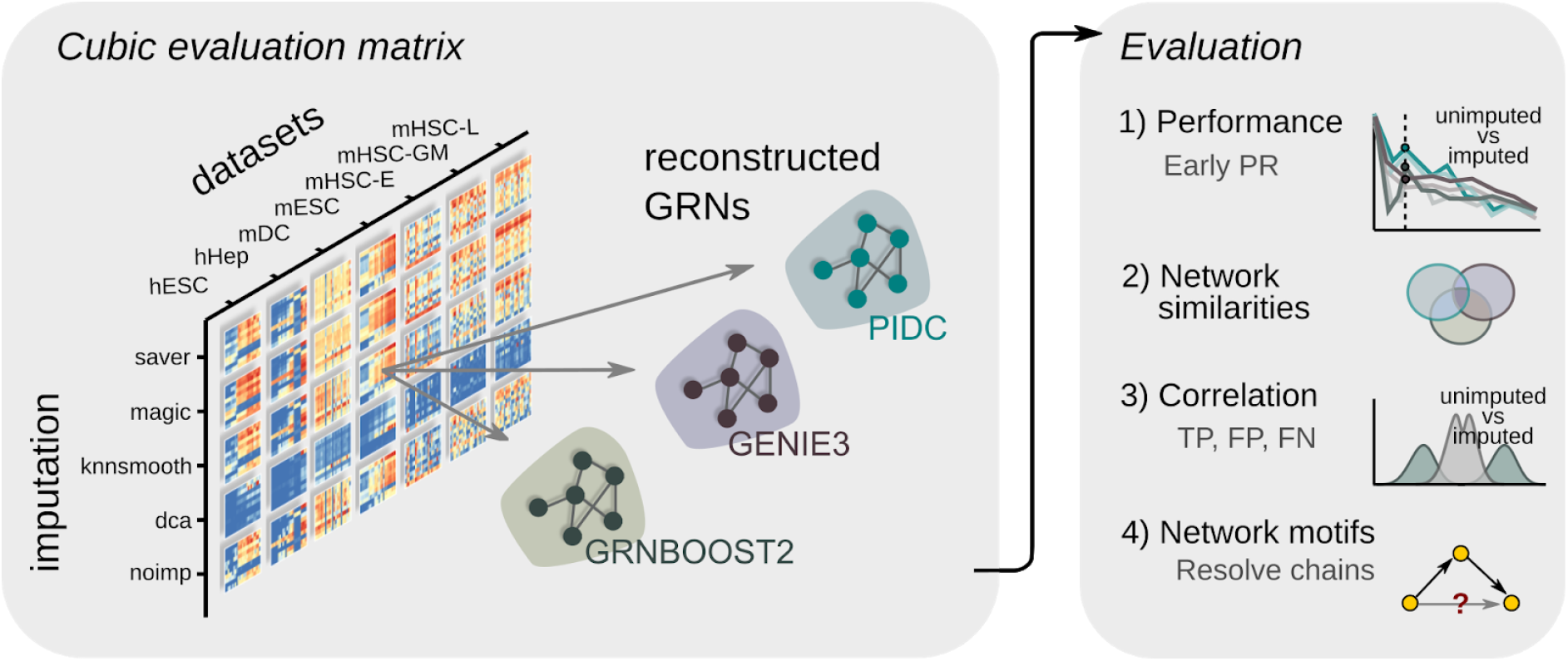
Systematic evaluation of network reconstruction from imputed and unimputed data. Cubic evaluation matrix consists of seven cell types from experimental scRNAseq data, four imputation methods (see text) and three network reconstruction algorithms. Imputed and unimputed (“noimp” in the Figure) scRNAseq data provide input expression matrices which are used by the gene regulatory network (GRN) reconstruction algorithms using the BEELINE framework (6). We evaluate the performances using the early precision ratios (EPR) and compare network results across different models. Additionally, we inspect the effect of gene-gene correlation on prediction classes (true positives (TP), false positives (FP), false negatives (FN)) before and after imputation, and we search for common motifs within the reconstructed networks. hESC: human embryonic stem cells, hHep: human hepatocytes, mDC: mouse dendritic cells, mESC: mouse embryonic stem cells, mHSC-E, mHSC-GM, mHSC-L: mouse hematopoietic stem cells - erythrocytes, granulo monocytes, lymphocytes.

Information on the seven cell types was derived from five experimental scRNAseq datasets: human embryonic stem cell (hESC) (14), human hepatocytes (hHep) (26), mouse embryonic stem cell (mESC) (27), mouse dendritic cells (mDC) (28) and mouse hematopoietic stem cells (mHSC) (29) that were further separated into the following subtypes: erythrocytes (mHSC-E), granulo monocytes (mHSC-GM) and lymphocytes (mHSC-L).

For the four imputation methods, we chose the following methods: two smoothing-based tools *magic* (18) and *knn-smoothing* (22); a model-based tool *saver* (17) and a deep-learning based tool *dca* (20). We included *dca* because the authors specifically expect to improve network reconstruction. A baseline model was established using normalized but unimputed data.

As for GRN reconstruction, we selected three tools: an information-based tool PIDC (4), and two tree-based tools GENIE3 (30) and GRNBoost2 (31). In the remainder of this paper we use the term “model” to refer to the combination of a GRN reconstruction algorithm with an imputation method or no imputation, respectively. We obtain the ground truth network from the STRING database — a functional protein-protein interaction network (32) and use the evaluation framework BEELINE (6) for measuring the performance of each network model (see Methods). Furthermore, we inspect the reconstructed network and compare the results with one another.

### 2.2 Imputation does not improve the performance of network reconstruction in general

A compact overview of the results obtained under all the models is provided in Figure 2, where each box summarizes results for one GRN reconstruction method. The performance measurements achieved by the respective model on the seven data sets are arranged on a vertical axis. Two performance measures have been computed: the early precision ratios (EPR) (6) which are shown in the three boxes of Fig. 2A, and the log_2_-ratios between epr_imputed_ and epr_unimputed_ which are shown in the three boxes of Fig. 2B. EPR refers to the number of true positive interactions within the top-k network normalized by the network density. Here, *k* refers to the number of positive interactions found in the ground-truth network (see Methods). An EPR of 1 indicates a random predictor. The second performance measure compares the performance of an imputation method relative to the performance of not using imputation. Here, a value of zero means no change, while a negative value indicates a detrimental effect of the imputation.

**Figure 2.**
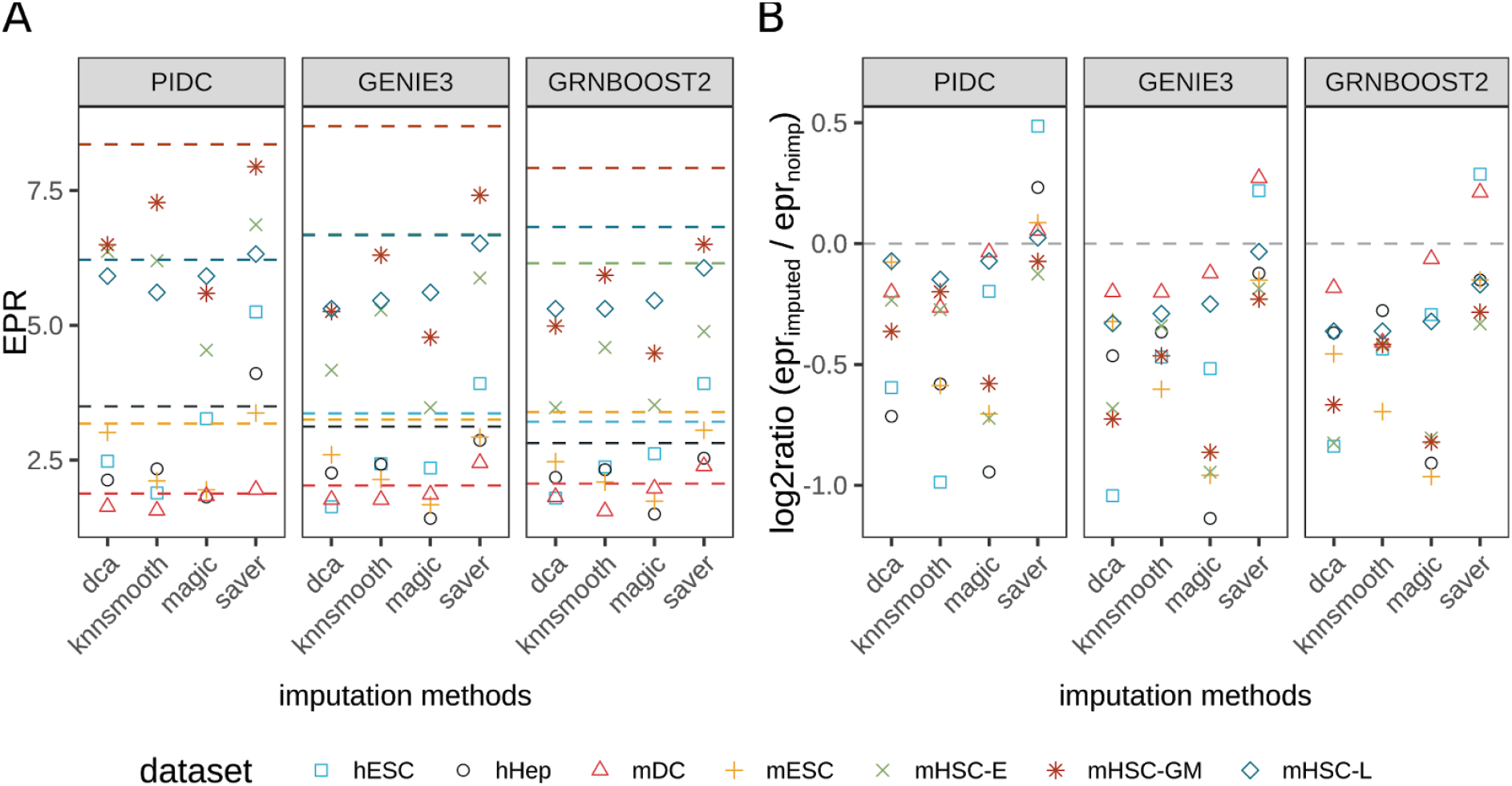
Impact of imputation on network reconstruction performances. (A) Absolute EPR scores across imputation methods (x axis label) and GRN inference algorithms (box) on seven different cell types (coded by shape and color). Dashed lines represent EPR scores obtained without imputation. EPR = 1 corresponds to a random predictor. (B) log2-ratios between EPR scores obtained using imputed and unimputed data. Log2-ratio = 0 represents no change in performance (grey dashed line) after imputation.

The EPR scores for unimputed data that were reported by Pratapa et. al. (6) could be reproduced in our analysis and are illustrated as a dashed line in Figure 2A. Results vary strongly with the datasets; the scores range from approximately 2 (for the mDC dataset) to 8 (for mHSC-GM), with less variation across GRN reconstruction algorithms. Applying imputation with either *dca*, *knnsmooth* or *magic*, does not improve the performance in any of the GRN reconstruction methods. While in mDC data the performance scores in each model scatter around the unimputed model, in mHSC-GM data the performance scores vary strongly, dropping from 8 to just below 5 for the *magic/GENIE3* model.

Focussing on the change of performance due to imputation as measured using the log2-ratios between imputed and unimputed EPR scores, we observe that only *saver* is able to improve the performance (Fig. 2B). The *saver*/PIDC model achieves log-fold-ratios up to +0.5 in 5 out of 7 datasets and 2 out of 7 datasets combined with GENIE3 or GRNBoost2. All other imputation methods worsen the performance with log-fold-ratios down to −1 which represents a performance decline of 100% in comparison to the unimputed model.

We further asked the question whether data quality as given by sequencing depth is a determinant of the success of imputation prior to GRN reconstruction. To answer this, we simulated cells *in silico* by downsampling the gene counts of the given experiments to 60% of their sequencing depth, thereby lowering the detection rate (Supp. Fig. 1). The hope would be that imputation has a more beneficial effect in these simulated data sets as compared to the original, higher quality data. However, similar results as above were obtained when we subjected the lower quality *in silico* data to our analysis pipeline (Supp. Fig. 2). Like with the original datasets, *saver/*PIDC obtain the highest improvements compared to the downsampled unimputed datasets. Nonetheless on downsampled data, *dca, knnsmooth* and *magic* are able to improve performance in some of the tested datasets, although not consistently.

Overall, our results demonstrate that our model performances are highly dataset-dependent. Applying imputation on the original data resulted mostly in a drop of performance of GRN reconstruction compared to the unimputed model, although potentially improving performance on low-quality data tested *in silico,*

### 2.3 Imputation method rather than GRN method determines results

The analysis presented in the preceding Section raises the question how strongly either the choice of imputation method or of network reconstruction algorithm affects the results. To answer this question we first address the variability in results when varying either the one or the other, and then study similarity among computed networks across the models.

With regard to the performance variability, we compare the variance of EPR log-fold-ratios under a fixed GRN reconstruction algorithm while varying across imputation methods, and, vice versa, varying the GRN algorithm while keeping the imputation method fixed. As Figure 3A shows EPR log-fold-ratios vary much more strongly when the GRN reconstruction algorithm is fixed than than the other way round (wilcoxon-test p-value ~7.86×10^-6^). This implies that the choice of imputation method determines the quality of results to a larger degree than the choice of GRN reconstruction algorithm.

**Figure 3.**
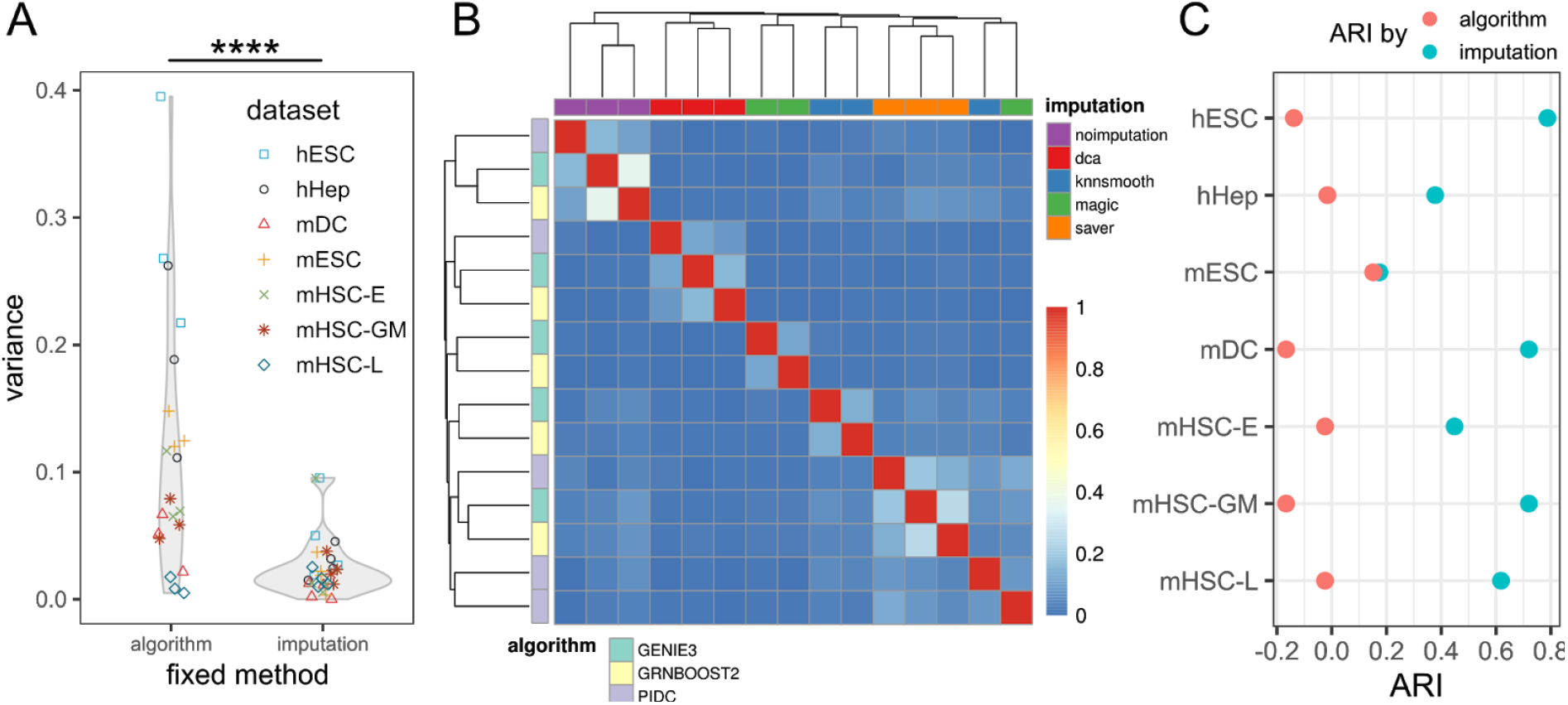
Variability in network results largely stems from imputation methods. (A) Variance distribution of EPR scores across imputation methods. Left violin plot keeps the GRN algorithm fixed and depicts the variances in EPR log-fold-ratios for each dataset across the imputation methods. Right violin plot shows the variances for fixed imputation methods. **** corresponds to p-value ≤ 0.0001 by wilcoxon rank sum test. (B) Clustered heatmap of network similarities measured by Jaccard index within top 500 reported interactions. Columns are color-coded by imputation methods. Rows are color-coded by network inference algorithms. (C) Adjusted rand index (ARI) obtained for clustering results in each cell type by annotation label “algorithm” (pink) and “imputation” (blue), respectively.

A direct consequence of this observation is the suspicion that the topology of the predicted networks may also be largely determined by the imputation method and to a lesser degree by the GRN reconstruction method. To test this, we inspect the overlap among the 500 most important gene-gene interactions of the computed networks. Here, we calculate pairwise similarity scores using the Jaccard index and use it to hierarchically cluster the networks. We found that networks tend to cluster with respect to imputation methods but not GRN methods (Fig. 3B, Supp. Fig. 4). To make this more precise, we use as a measure of cluster purity the adjusted rand index (ARI) (33,34). ARI coefficients calculated across the seven different cell types show higher cluster purity when labelled with imputation method as opposed to network reconstruction algorithms (Fig. 3C).

**Figure 4.**
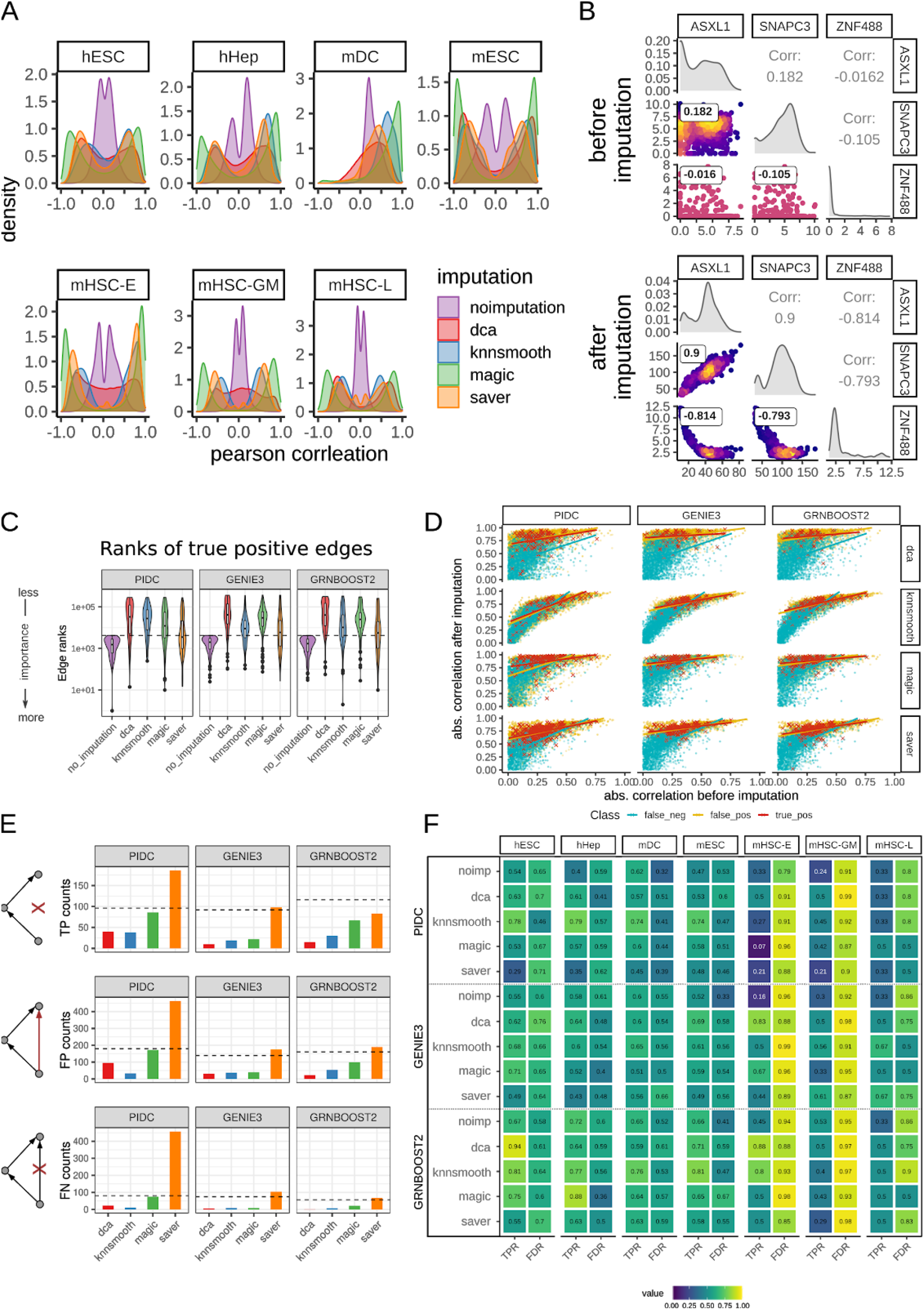
Gene-gene correlation before and after imputation and its impact on the predicted interactions. (A) Gene-gene correlation distributions obtained in each cell type color-coded by imputation method among top 500 most variable genes and significantly varying TFs. (B) Paired density scatter plots before and after imputation with *dca*. GRNBoost2 reported the pairwise interactions between ASXL1, SNAPC3 and ZNF488 among the top-k network after imputation in hESC data. (C) Change of edge ranks in true positive (TP) interactions identified by unimputed model after imputation in hESC data. Dashed line indicates the rank threshold corresponding to the top-k network. Interactions below the dashed line represent TP within the respective model. Low edge ranks represent highly important interactions. (D) Change of correlation values for TP (red crosses), FP (yellow dots) and FN (blue dots) classified by each model in hESC data. Positively predicted interactions differ clearly from FN interactions. (E) Counts of positively and negatively predicted network chain motifs in hESC data for each model. TP network chains agree both in prediction and ground truth. FP network chains are falsely positively predicted chains being actual feed-forward loops in the ground truth. FN network chains are falsely predicted as being feed-forward loops when they are actually network chains in the ground-truth network. (F) TPR and FDR scores for network chain motifs obtained by statistics in E). Ideally, TPR values should be close to 1 whereas FDR values should be close to 0.

We conclude that the imputation method largely determines model performance, leaving little influence to the subsequent GRN reconstruction algorithm. The choice of imputation method further biases the outcoming network leading to little consensus across the most important recovered gene-gene interactions as computed based on different imputation methods.

### 2.4 Inflation of gene-gene correlations and its impact on the network topology

Based on the reported results, we examine how imputation generally affects gene-gene correlation coefficients. Although not all network reconstruction algorithms use correlation-based measures to recover interactions, we still use Pearson’s correlation coefficient as a proxy for the association between two genes. Subsequently, we will investigate whether the interactions within the reconstructed networks affect the global network structure.

Exploring the overall distributions of gene-gene correlations after imputation on scRNAseq data we observe a strong increase in gene-gene correlations (Fig. 4A). Generally, gene-gene correlations go from almost no correlation when computed using unimputed data to very good anti- and positive correlations due to imputation. Here, *magic* l eads to the most extreme enhancement. More specifically, Figure 4B exemplifies the association between three genes before and after imputation, transforming very weak correlations to almost perfect (anti-)correlations. These associations were only reported after imputation using *dca* among the top-k network using GRNBoost2 in hESC data. Indeed, we commonly find such associations across different datasets and imputation methods.

In order to see what impact this enhancement of correlation has on the network structure we next investigated the network density after imputation in relation to the unimputed data using log-ratios (Supp. Fig. 3A). Here, we looked at the top-k networks according to the EPR score. Imputation methods alter the network densities with log-ratios ranging from −0.5 and +0.5 in hESC, hHep, mDC and mESC data, except for *saver* and PIDC in hESC data with a slightly higher value of 0.59. For the three subtypes of mHSC data we observe larger changes in network density reaching log-ratios beyond ±1. Especially here, imputations combined with GENIE3 and GRNBoost2 lead to a sparser network whereas all combinations of imputation methods with PIDC lead to a denser network structure. This is not surprising as GENIE3 and GRNboost2 are network reconstruction algorithms that provide a directed network, whereas PIDC results in undirected interactions providing a backward and forward edge with the same ranks. As we select top-k ranked interactions we take interactions sharing the same ranks simultaneously, consequently leading to a denser network with PIDC.

Besides network density, the network topology is also determined by the node degree distribution. Before imputation we observe a heavy tail node degree distribution predominantly in GENIE3 and GRNBoost2 indicating the presence of many hub nodes (Supp. Fig. 3B). After imputation the heavy tail disappears when using *dca, magic* and *knnsmooth* while it still exists when using *saver.* Generally, PIDC does not lead to this structural change in node degree distribution.

As a conclusion, the enhancement of gene-gene correlations due to imputation appears to lead to notable changes in the topology of the predicted gene networks.

### 2.5 Increased correlation values in false positive interactions inflate network results

Since we have observed that imputation may decrease the performance of GRN network reconstruction, we attempt to understand how the altered correlations in imputed data affect network reconstruction. To this end, we explore the change of edge ranks and correlation values of the reported (i.e., positively predicted) and missed (i.e., negatively predicted) interactions.

Overall, the ranks of true positive (TP) interactions reported in the unimputed data change significantly after imputation (Fig. 4C, Supp. Tab. 1, Supp. Fig. 5). Some of the previously reported TP interactions could be recovered after imputation. Nevertheless, the majority of previously reported TP interactions shift after imputation towards the end of the gene interaction ranking list, and are considered less important. As a consequence, other interactions become more important.

Therefore, we look at the change of correlation of positively predicted interactions before and after imputation. Figure 4D (and Supp. Fig. 6) show scatter plots of gene-gene interactions with the absolute values of correlation coefficients before imputation on the horizontal axis and the correlation coefficient after imputation on the vertical axis. For each model, red dots are the true positive interactions, yellow are the false positives, and blue are the false negatives. The general shape of the scatter plot reiterates the observation that correlation coefficients tend to get enhanced by imputation. For each class we computed regression lines. For better recognition of true positives after imputation, one would hope for the TP regression line (shown in red in Fig. 4D) to lie well above the others - which is not really the case. We generally observe a strong enhancement of correlations as indicated by the height of the intercept of the regression lines. In 11 out of 12 cases the regression lines for both true and false positive predictions are almost congruent with each other. Note that the red color dominates the other ones and the dots below a red one are not visible.

Interestingly, we see remarkably different regression lines if we take the false negative (FN, blue) interactions into account. The majority of FN correlations remain low after imputation, as indicated by the height of the intercept in Fig. 4D. Presumably, the FN correlation values that actually get enhanced get lost in the background due to the inflation of FP correlations in the inferred top-k network. Thus, the boost of correlation values makes it harder for GRN reconstruction methods to separate the actual signal from the background.

Many GRN reconstruction methods have the goal of distinguishing direct interactions from transitively inferred ones (35). Therefore, we tested whether the GRN reconstruction algorithms analyzed in this study are able to make the necessary distinction. Given three genes X, Y, and Z where X is correlated with Y, and Y is correlated with Z, these genes form a network chain. However, oftentimes by transitivity these associations seem to imply a correlation between X and Z, thus forming a network loop. Generally, in network theory it is challenging to distinguish chains from loops. In this context, we analyze how the models deal with the identification of network chains from imputed data. Errors are counted if a supposedly false loop is detected or a chain ist detected instead of a loop (Fig. 4E).

In general, PIDC identifies the highest number of network motifs independent of the imputation used. Using *saver* in combination with PIDC one is able to find the highest number of TP chains. However, *saver*/PIDC mistakenly identifies network motifs at the same time. In order to measure the performance between true and false predictions we calculate the true positive rates (TPR) and false discovery rates (FDR) for each network inference and imputation method applied to each dataset (Fig. 4F).

The performance of network motif search among the top-k networks does not seem to be affected by imputation. Hence, either imputation methods do not necessarily induce transitive correlations or the network reconstruction methods deal well with transitively induced correlations.

## 3 Discussion

The advent of single-cell transcriptomics has rekindled the interest in reconstructing gene regulatory networks from transcriptomics data, primarily for two reasons. Firstly, it is of great interest to study regulation in individual cells in the hope to eventually uncover how, e.g., differentiation processes proceed. Secondly, the main obstacle in gene network reconstruction from bulk transcriptome data appears to be the low number of available samples in comparison to the large numbers of genes. For example, simulations have demonstrated that high quality reconstruction of gene networks requires a much larger number of samples than the number of genes (35). Seeing each single cell as a sample, the expectation arose that single-cell transcriptomics would solve this conundrum by providing a sufficiently large number of samples, thus putting high quality network reconstruction within reach.

It was sobering for us to see that due to the sparse nature of single-cell RNA-seq data, individual cells cannot contribute as much information to network reconstruction as bulk samples. Indeed, preprocessing of single-cell data for data analysis is crucial (9), and is implemented in many computational pipelines. Imputation has become a possible element of this preprocessing in the hope it would supplement the missing information. In this study we have however demonstrated that the choice of imputation prior to GRN reconstruction influences the results in a two-fold manner: First, it affects the performance of network reconstruction leading to highly variable accuracies and, secondly, the reconstructed network differs significantly between imputation methods.

We have systematically evaluated the effect of imputation on GRN reconstruction using experimental scRNAseq data on seven cell types. In this context, we have demonstrated that overall, imputation does not lead to an improvement of GRN reconstruction. However, *saver* i n combination with PIDC may lead in some datasets to an increase in performance. We have shown and thereby agree with previous studies that imputation may boost gene-gene correlations in a dubious way, thereby introducing false positives (25). In turn, these false positives predispose network structures toward forming circular dependencies. In fact, if network reconstruction methods rely on associations that use correlation to some extent (for example regression-based methods) the circularity is highly redundant. Andrews et. al. have warned of this circularity before, albeit in the context of differential expression analysis (36). Consistent with our findings Andrews et. al. showed that *saver* introduces the smallest number of spurious gene-gene correlations. We speculate that the combination of *saver*/PIDC works well because *saver* is a model-based imputation method and PIDC is a mutual-information based algorithm; the two approaches follow independent assumptions complementing one another, thus avoiding the use of redundant information.

In this study we have tested our hypothesis on experimental datasets with fairly large library sizes and gene detection rates (Supp. Fig. 1.). In order to test our hypothesis on more shallowly sequenced single-cell experiments we lowered the detection rate introducing more zero counts *in silico*, Our results have shown that using *saver* with PIDC improves results in most cases. However, generally we discourage the indiscriminate usage of imputation prior to GRN reconstruction because imputation tends to introduce a bias into the derived networks. If need be we recommend the use of *saver* and PIDC. It should be noted that we are not discouraging imputation in general. There may be many other applications that are not studied here, where imputation can be useful, depending on the type of analysis that is subsequently performed.

## 4 Methods

### 4.1 Data collection and preprocessing of scRNAseq data

We collected preprocessed and normalized experimental scRNAseq count data provided in the BEELINE paper (6). Here, the authors also provide the corresponding pseudotime for each dataset / cell type. Please refer to the BEELINE paper for information about preprocessing, normalization, and pseudotime inference.

However, *dca* needs unnormalized raw count data. Therefore, we downloaded the fastq files using the corresponding accession numbers: GSE75748 (hESC) (14), GSE81252 (hHEP) (26), GSE98664 (mESC) (27), GSE48968 (mDC) (28) and GSE81682 (mHSC) (29). For human and mouse we aligned the fastq files to hg19 (GENCODE release 29) or mm10 (GENCODE release M19), respectively and counted the reads per gene using STAR (version 2.7.4a) (37).

Following the BEELINE approach, using normalized count data we select the top 500 most variable genes across pseudotime using a general additive model (‘gam’ R package). In addition to these genes we also include significantly varying TFs (Bonferroni corrected p-value < 0.01).

We filter both imputed and unimputed scRNAseq data using the same set of (i) top 500 most variable genes and (ii) all significantly varying TFs, in order to make a fair comparison between networks inferred using imputed and unimputed data.

### 4.2 Code availability

All relevant scripts and R notebooks for reproducing the results are available at Github (https://github.com/lylamha/imputation_GRNinference). The release includes tutorials from data imputation to the evaluation of the reconstructed networks. It covers the evaluation pipeline with the corresponding analyses and plotting results.

### 4.3 Imputation

To impute scRNAseq data we use *dca* (version 0.2.3), *knnsmooth* (version 2.1), *magic*(‘Rmagic’ R package version 2.0.3) and *saver* (‘SAVER’ R package version 1.1.2). Our rationale for selecting *knnsmooth*, *magic* and *saver* is based on a previous comprehensive benchmark evaluation of various imputation methods (23). Additionally, we also include *dca* as it has been explicitly recommended as improving GRN reconstruction.

We apply each imputation method to normalized count data except for dca where we use the raw counts:

~~~
dca </path/to/ExpressionData_raw.csv></path/to/dca_result_folder>
python3 knn_smooth.py -k 15 -d 10 \
-f <path/to/ExpressionData.csv> \
-o <path/to/ExpressionData_knnsmooth_imputed.csv> --sep,
magic Rsnippet:
# so_dat (seurat object using library(Seurat))
so_dat <- magic(so_dat, assay=“RNA”, genes=“all_genes”)
Dat.magic <- as.data.frame(so_dat@assays$MAGIC_RNA@data)
saver Rsnippet:
saver_res <- saver(input_expr_matrix, size.factor = 1, ncores = 20,
estimates.only= F)
dat.saver <- as.data.frame(saver_res$estimate)
~~~

### 4.4 Network reconstruction via BEELINE

Several tools have been developed to infer GRNs from scRNAseq data differing in their algorithmic approach. They can be categorized into four main classes: correlation-, regression-, mutual information- or modelling-based approaches (6). In this study we evaluated PIDC, GENIE3 and GRNBoost2 which have been previously recommended by Pratapa et. al. (6). We use the imputed and unimputed scRNAseq data as input matrices for network reconstruction with PIDC, GENIE3 and GRNBoost2 using default parameters. To this end, we use the evaluation framework BEELINE (version 1.0).

As part of the BEELINE pipeline we first run ‘BLRunner.py’ to reconstruct the networks. Then, we filter the reconstructed networks in order to only include interactions from TFs to genes.

Finally, we use ‘BLevaluater.py’ to compute early precision scores evaluating the performance of each network by comparing it to a ground truth network. Here, we choose the functional protein-protein interaction database STRING and filter for genes that only occur in the input expression matrix.

By using early precision scores we only analyze the top-k networks.

### 4.5 Characterizing the reconstructed networks

#### 4.5.1 Top-k network

For comparability reasons we focus our analyses on the top-k networks. The top-k network of a reconstructed network includes the first k interactions selected by their ranks which were assigned by descendingly ordered edge weights. Here, *k* represents the number of positive interactions in the ground truth network. Interactions can share the same ranks, e.g., the forward and backward interactions in an undirected graph. So with *k* interactions reported in the ground truth network we select all interactions which ranks are lower than or equal to *k* obtaining the top-k network. Note, that the number of reported interactions can be higher than *k.*

#### 4.5.2 Network density and node degree

Taking into account the interaction between transcription factors and genes only the network density is calculated by numEdges / ((numGenes * numTFs) - numTFs).

In order to calculate the node degree we consider all out- and incoming edges for a given node.

### 4.6 Methodology of evaluation

#### 4.6.1 Early Precision Ratios (EPR)

We evaluate the performance of each inferred network based on using early precision sores (EP) which is given by the number of TP divided by the number of positively predicted observations within the top-k network. EP scores were calculated using BEELINE. Each dataset has a different underlying ground truth subnetwork, hence different evaluations regarding the random predictor. To account for these differences and in order to maintain comparability across datasets we divide the EP scores by the network density (see formula above) of each ground truth subnetwork obtaining EP ratios (EPR). Thus, EPR of 1 is indicative of a random predictor in all experimental datasets. To compare the performance of network inference in each imputation method with the corresponding unimputed data, we calculate log2-ratios between EPR_imputed_ and EPR_unimputed_.

#### 4.6.2 Network similarities

In order to compare similarities across the reconstructed networks we select the top 500 interactions reported in each model. Given two networks, similarity scores are obtained by the Jaccard index which is defined as the number of overlapping interactions divided by the number of unified reported interactions. Repeating this in a pairwise iterative manner we obtain a similarity matrix which we use as an input for a heatmap that is clustered row- and column-wise (‘pheatmap’ R Package).

We calculate adjusted rand index (ARI) scores (‘mclust’ R package) in order to evaluate the clustering results based on an annotation label (34). As annotation labels we use the network reconstruction algorithm as well as the imputation method. We compare ARI scores across datasets obtained by the two labels using the pairwise wilcoxon rank sum test.

## Acknowledgements

We thank Mahsa Ghanbari and Prabhav Kalaghatgi for comments and proofreading our manuscript, Stefan Börno for assistance with the single-cell RNAseq processing, and Thomas Kreitler for technical support.

## Author information

### Contributions

L.H.L. and M.V. designed the study. L.H.L processed and analyzed the data. L.H.L and M.V. wrote the manuscript.

### Corresponding author

Correspondence to Martin Vingron

