## Supplementary figures and table for "Effect of imputation on gene network reconstruction from single-cell RNA-seq data"

### original scRNAseq dataset

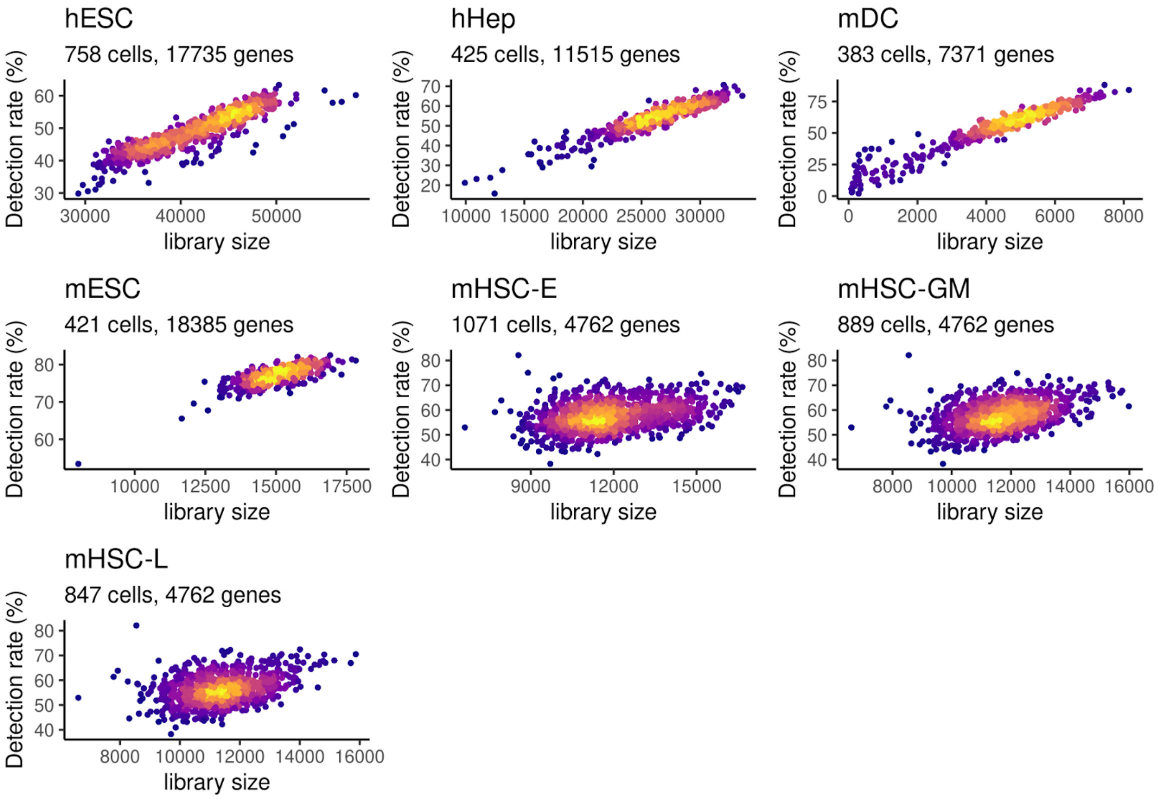


### downsampled scRNAseq data (60% of sequencing depth)

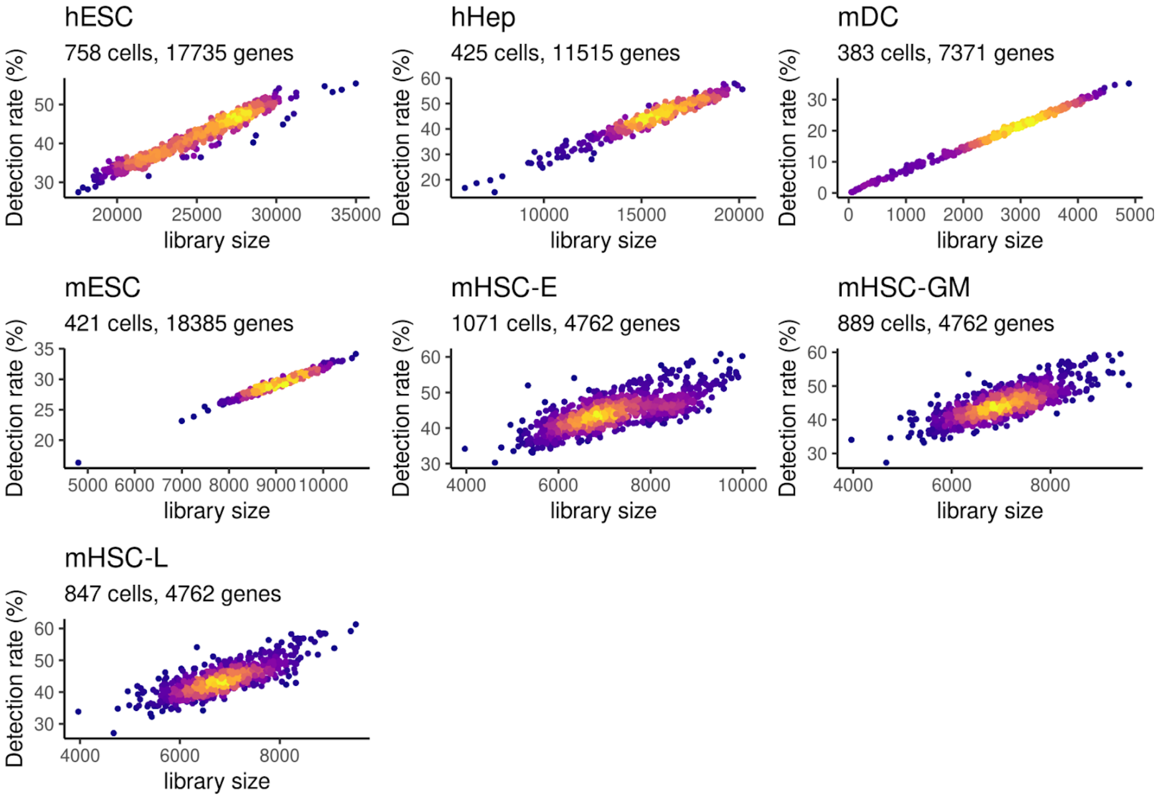

**Supp. Fig. 1: Gene detection rate and library size in experimental scRNAseq datasets (original and downsampled).** Scatterplots colored by density of points (cells). Gene detection using a threshold of gene count > 0. Library size determined by the sum of all gene counts. Downsampling procedure performed by sampling n times (60% of the original library size) according to the multinomial distribution.

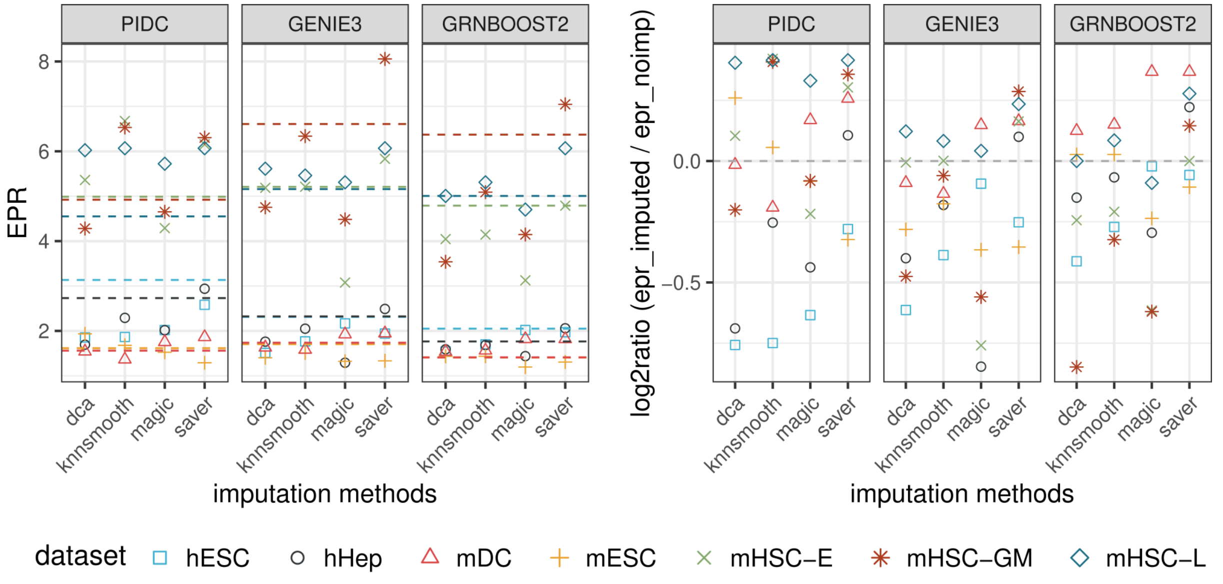


**Sup. Fig. 2: Performance measures of network models obtained by downsampled dataset.** (Left) Similar representation of performances as in Figure 2. Absolute EPR scores on downsampled scRNAseq data (60% of original library size). Dashed line represents EPR scores obtained without imputation. (Right) log2-ratios between imputed and unimputed EPR scores. Log2ratio = 0 represents no change in performance (grey dashed line) after imputation. More improvements (positive log2ratios) than in Figure 2.

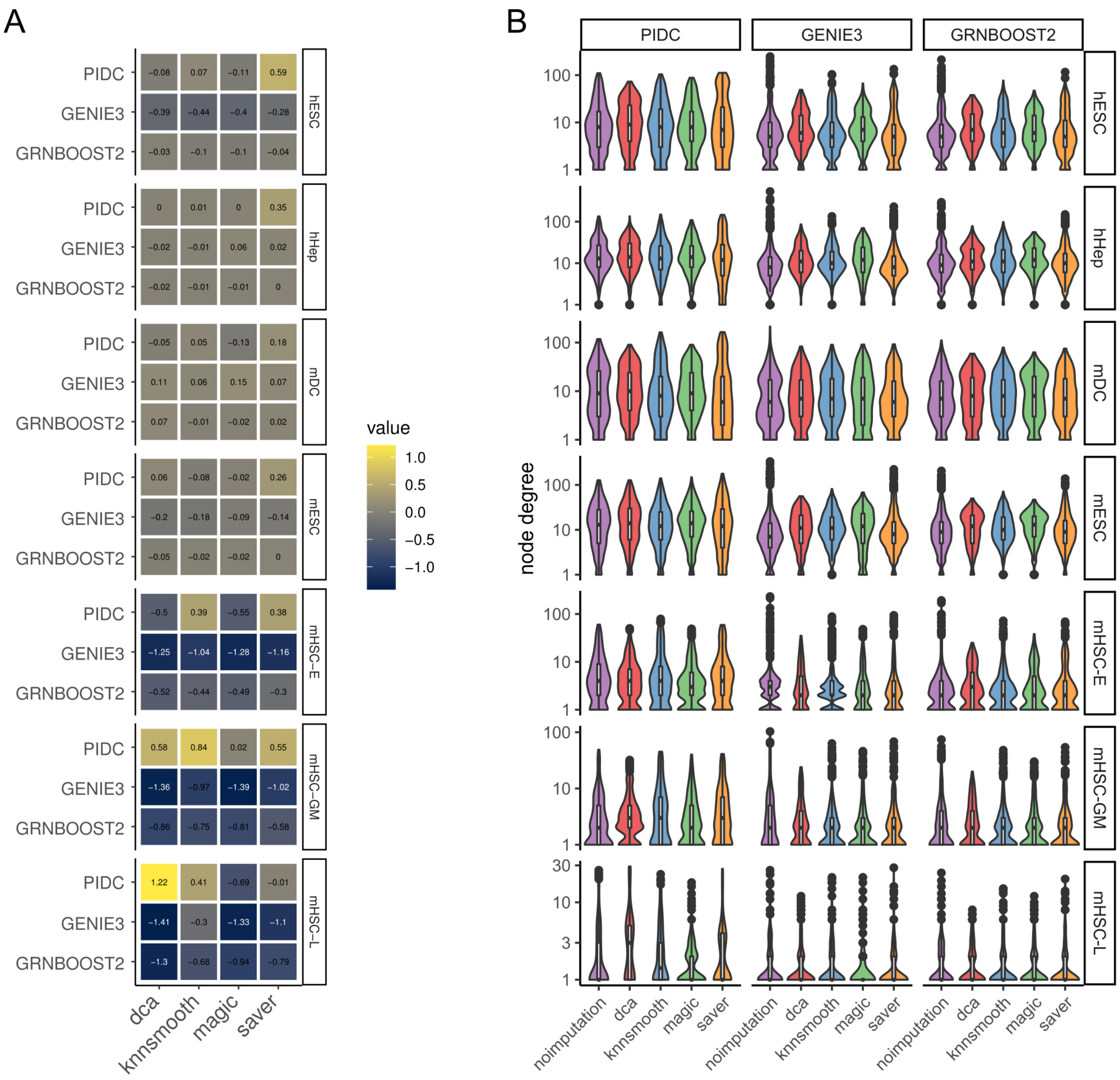


**Supp. Figure 3 | Structural changes in inferred networks.** (A) Change of node density before and after imputation. Log2 ratios between density (after imputation) and density (before imputation) are color-coded. Positive values represent a denser whereas negative values represent a more sparse network with respect to the unimputed model. (B) Node degree distribution across all models. Y-axis is log-scaled.

| data | imputation | PIDC | GENIE3 | GRNBOOST2 |
|---|---|---|---|---|
| hESC | dca | 6.23E-59 | 1.35E-52 | 3.22E-57 |
| | knnsmooth | 9.30E-70 | 5.56E-30 | 1.72E-35 |
| | magic | 5.41E-45 | 1.89E-47 | 1.68E-51 |
| | saver | 1.99E-15 | 5.46E-12 | 8.93E-13 |
| hHep | dca | 2.77E-144 | 3.67E-89 | 5.47E-83 |
| | knnsmooth | 1.11E-130 | 3.88E-85 | 5.87E-86 |
| | magic | 1.26E-126 | 1.19E-125 | 1.86E-108 |
| | saver | 1.66E-23 | 1.08E-37 | 1.34E-29 |
| mDC | dca | 1.15E-48 | 1.91E-39 | 8.93E-42 |
| | knnsmooth | 1.47E-46 | 5.11E-38 | 3.20E-38 |
| | magic | 2.14E-58 | 6.35E-31 | 9.12E-32 |
| | saver | 2.69E-10 | 7.78E-08 | 3.66E-10 |
| mESC | dca | 2.85E-75 | 6.53E-79 | 7.09E-83 |
| | knnsmooth | 2.84E-75 | 9.74E-59 | 5.33E-77 |
| | magic | 1.54E-78 | 1.25E-80 | 4.75E-90 |
| | saver | 3.61E-05 | 3.28E-07 | 7.37E-10 |
| mHSC-E | dca | 1.93E-08 | 2.97E-24 | 6.55E-25 |
| | knnsmooth | 1 | 4.70E-07 | 1.72E-10 |
| | magic | 2.44E-14 | 6.80E-27 | 4.39E-26 |
| | saver | 1 | 0.001687 | 0.0004226 |
| mHSC-GM | dca | 5.32E-44 | 1.02E-33 | 7.75E-32 |

| | | | | |
|---|---|---|---|---|
| | knnsmooth | 0.05846 | 2.32E-09 | 9.27E-17 |
| | magic | 8.66E-21 | 2.31E-23 | 2.89E-22 |
| | saver | 1 | 8.45E-07 | 6.18E-09 |
| mHSC-L | dca | 1 | 0.006637 | 1.51E-05 |
| | knnsmooth | 1 | 1 | 0.3427 |
| | magic | 0.005388 | 0.004132 | 3.54E-06 |
| | saver | 0.7522 | 1 | 0.0006845 |

**Suppl. Tab. 1:** Corrected p-values (Bonferroni method) obtained after Wilcoxon rank sum test between ranks of unimputed true positive edges and their respective ranks after imputation.

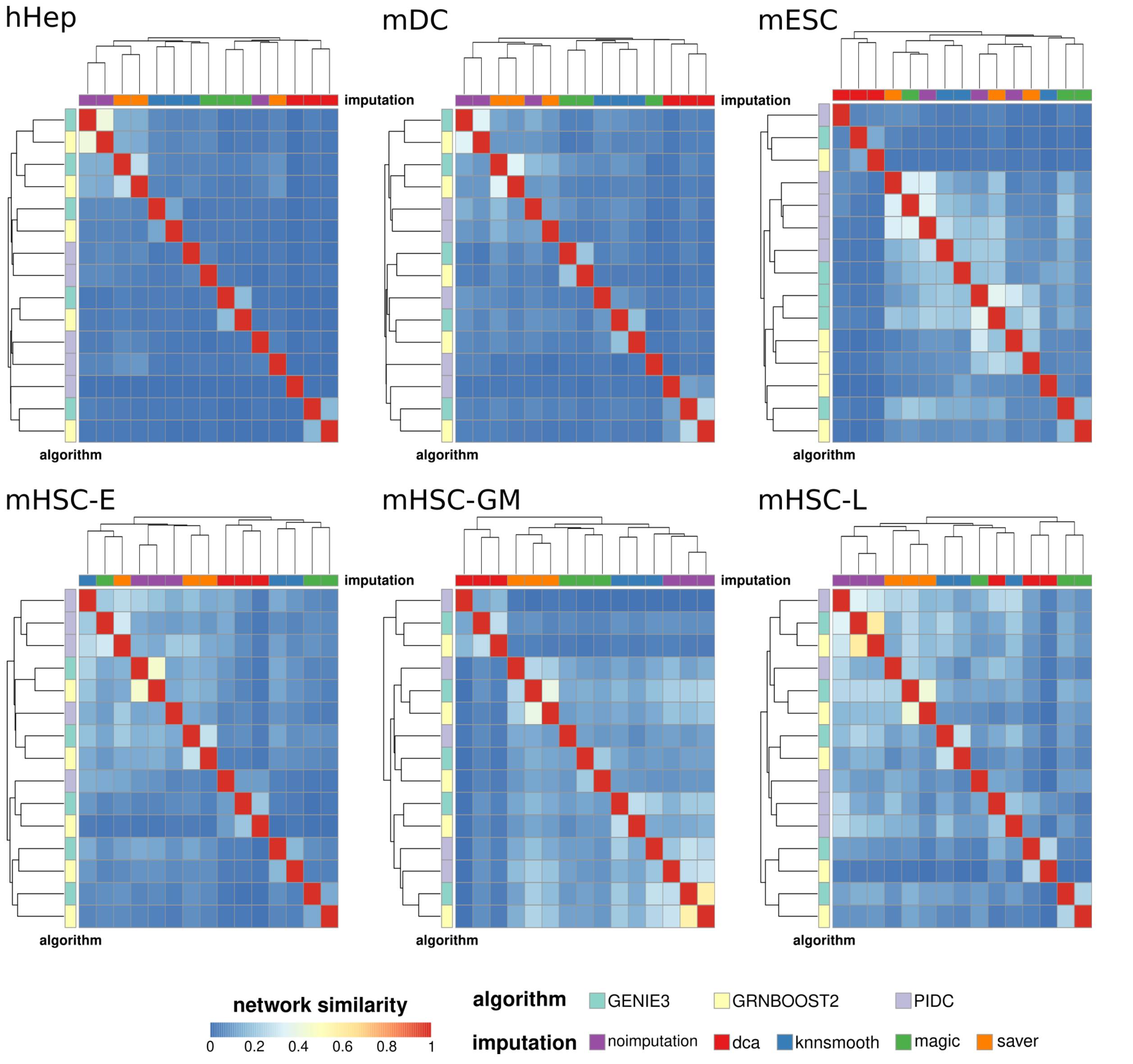


**Supp. Fig. 4: Network similarities across all models and cell types.** According to Figure 3B we inspect the heatmap of network similarities of the remaining cell types. Network similarity scores obtained by pairwise Jaccard index from top500 interactions. Columns are annotated by imputation method, rows are annotated by GRN reconstruction algorithm.

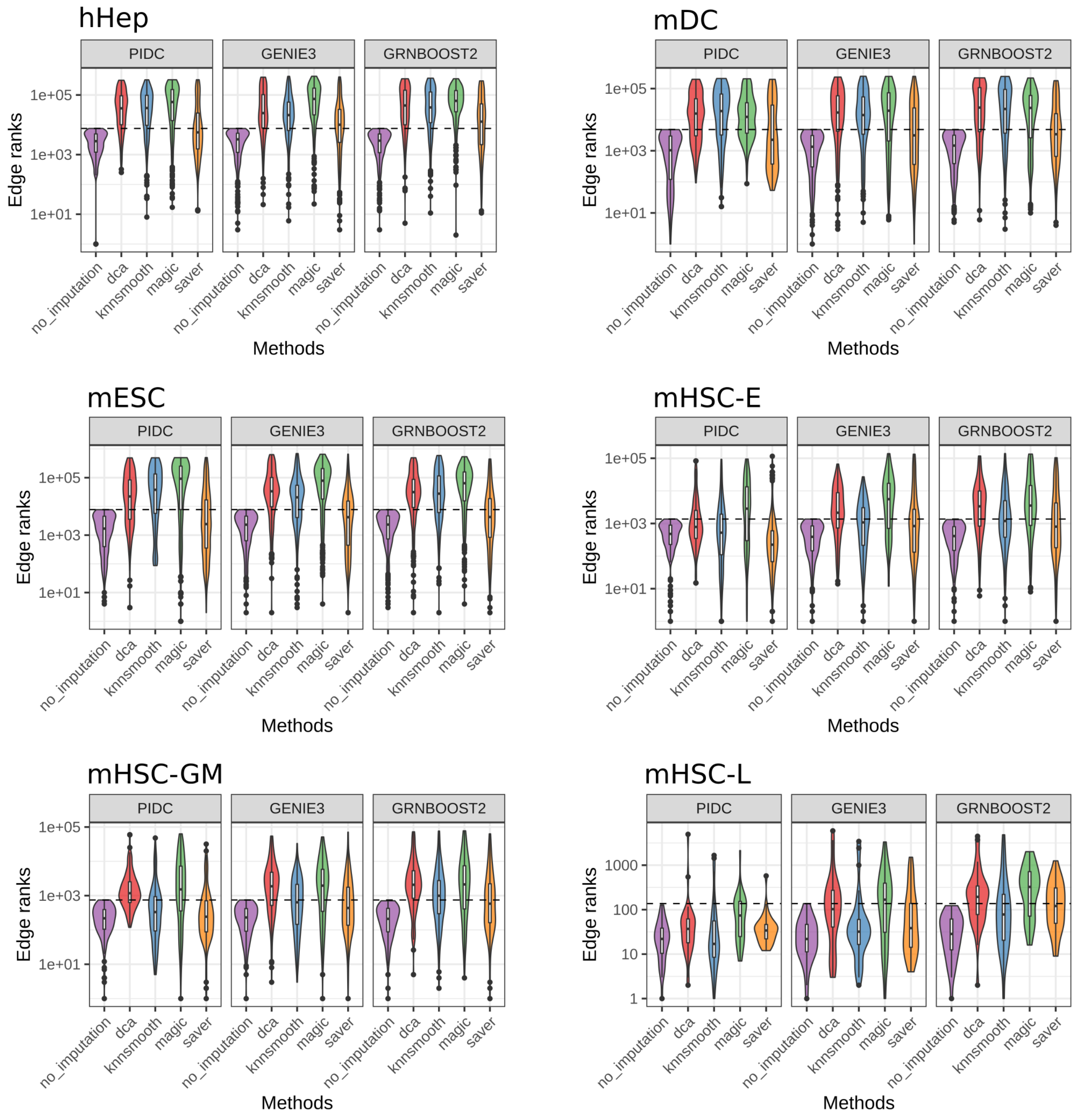


**Supp. Fig. 5: True positive interactions identified on unimputed data and their change in edge ranks after imputation.** According to Figure 4C we inspect the change of unimputed TP ranks after imputation in the remaining cell types. Corrected p-values obtained by Wilcoxon rank sum test can be taken from Suppl. Tab. 1.

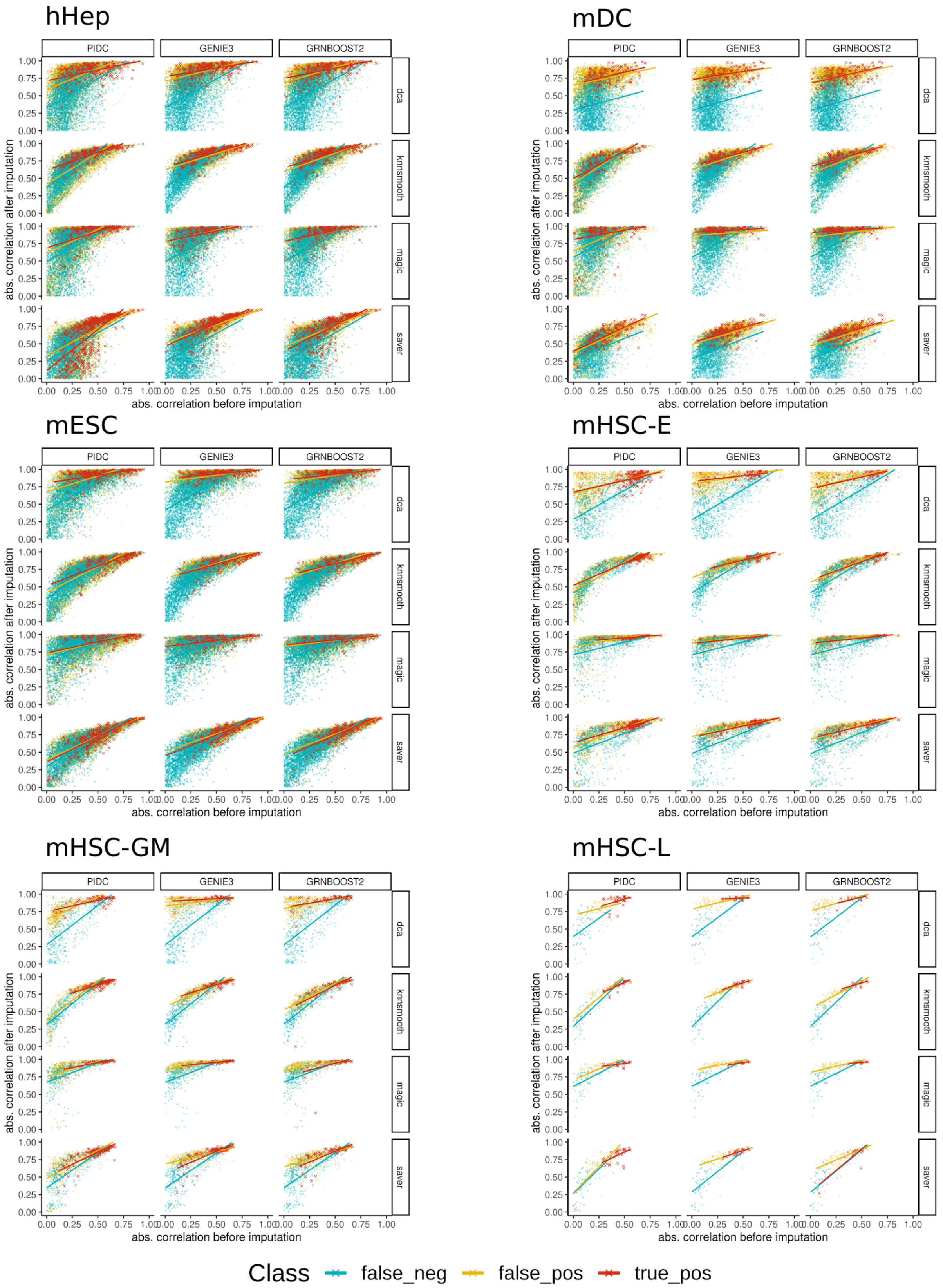
hHep
mDC
mESC
mHSC-E
mHSC-GM
mHSC-L
PIDC
GENIE3
GRNBOOST2
dca
knnsmooth
magic
saver
0.00
0.25
0.50
0.75
1.00
abs. correlation after imputation
abs. correlation before imputation
Class
false_neg
false_pos
true_pos

**Supp. Fig. 6: Absolute Pearson's correlation coefficients before and after imputation colored by prediction class obtained in each model.** According to Figure 4D we inspect change of correlation values for TP, FP and FN classified by each model in the remaining cell types. Colors correspond to the prediction classes in each model.
